# Within-Individual Organization of the Human Cognitive Cerebellum: Evidence for Closely Juxtaposed, Functionally Specialized Regions

**DOI:** 10.1101/2023.12.18.572062

**Authors:** Noam Saadon-Grosman, Jingnan Du, Heather L. Kosakowski, Peter A. Angeli, Lauren M. DiNicola, Mark C. Eldaief, Randy L. Buckner

## Abstract

The human cerebellum possesses multiple regions linked to cerebral association cortex. Here we mapped the cerebellum using precision functional MRI within individual participants (N=15), first estimating regions using connectivity and then prospectively testing functional properties using independent task data. Network estimates in all participants revealed a Crus I / II cerebellar megacluster of five higher-order association networks often with multiple, discontinuous regions for the same network. Seed regions placed within the megaclusters, including the disjointed regions, yielded spatially selective networks in the cerebral cortex. Compelling evidence for functional specialization within the cerebellar megaclusters emerged from the task responses. Reflecting functional distinctions found in the cerebrum, domain-flexible cerebellar regions involved in cognitive control dissociated from distinct domain-specialized regions with differential responses to language, social, and spatial / episodic task demands. These findings provide a clear demonstration that the cerebellum encompasses multiple zones dedicated to cognition, featuring juxtaposed regions specialized for distinct processing domains.

## Introduction

Growing evidence suggests that the cerebellum contributes to non-motor functions and further that multiple regions associated with diverse cognitive and affective domains populate the primate cerebellar cortex, in particular the large region centered on and surrounding the Crus I and II lobules (see Leiner, Leiner, & Dow 1986; Schmahmann 1991, 2019; Strick et al. 2009; Stoodley & Schmahmann 2009; Buckner 2013; Keren-Happuch et al. 2014; Koziol et al. 2014; Sokolov, Miall, & Ivry 2017; Guell et al. 2018; King et al. 2019; Guell & Schmahmann 2020; Habas 2021). The present work explored the detailed functional organization of the cognitive domains of the human cerebellum using within-individual precision mapping methods combined with intensive study of function.

Characterizing the organization of the cerebellum confronts specific challenges due its distinct anatomical circuitry and its small size. First, anatomical connectivity between the cerebrum and cerebellum is polysynaptic, meaning that traditional tracer injection techniques cannot map the spatial relations between cerebellar and cerebral cortices. In studies using viral multi-synaptic tracing techniques, Strick and colleagues (Middleton & Strick 1994, 2001; Kelly & Strick 2003; see also Strick et al. 2009) demonstrated closed-loop circuitry between the Crus I / II lobules, which fall between the motor zones, and regions of prefrontal cortex (PFC). While these methods established an anatomical substrate for cerebellar contributions to cognition, there is no equivalent direct anatomical technique in humans.

The second challenge lies in the expected tight juxtaposition of cognitive domains in the cerebellum. Specifically, the organization of networks supporting cognitive functions has been difficult to disentangle in the cerebral cortex, especially within the association zones where there are side-by-side juxtapositions of small, functionally distinct regions that are blurred in methods that use group-averaging (Fedorenko et al. 2012; Laumann et al. 2015; Braga & Buckner 2017). The smaller cerebellum possesses an intricate and complicated surface geometry that presents an even greater challenge (Diedrichsen et al. 2009; Sereno et al. 2020). If the organization of the cerebellum parallels that of the cerebral cortex, there will be closely juxtaposed regions with idiosyncratic anatomical differences and spatial positions that vary from person to person (see Marek et al. 2018).

To overcome these challenges, the present work uses precision mapping fully within the individual to explore the detailed organization of the human cerebellum. Two complimentary techniques were combined -- resting-state functional connectivity MRI (fcMRI) and task-based functional MRI (fMRI). fcMRI provides a means to detect a proxy for anatomical connectivity between the cerebrum and cerebellum, and map candidate cerebellar regions coupled to distinct cerebral networks. Supporting the validity of the approach, fcMRI studies have demonstrated contralateral coupling between motor cortex and the expected regions of the cerebellum (Krienen & Buckner 2009) that is disrupted by lesions to the pons (Lu et al. 2011). Moreover, distinct motor effector regions in the cerebellum can be mapped using fcMRI indicating that the technique has spatial specificity (Buckner et al. 2011; Guell et al. 2018). However, fcMRI does not assess function. To assess functional selectivity, task-based fMRI responses were measured within the same intensively studied individuals whose cerebellums were mapped. In this manner, multiple closely juxtaposed regions of the cerebellum could be delineated within the idiosyncratic anatomy of the individual and then the functional properties explored in independent task data.

Prior discoveries provide the foundation for our efforts. Building from the first finding of a cognitive response in the cerebellum using positron emission topography (PET; Petersen et al. 1989), there has been progress from both task-based fMRI (Desmond & Fiez 1998; Stoodley & Schmahmann 2009; Guell & Schmahmann 2020; King et al. 2019) and network analyses using fcMRI (Allen et al. 2005; Habas et al. 2009; Krienen & Buckner 2009; O’Reilly et al. 2010; Guell et al. 2018; Marek et al. 2018; Xue et al. 2021; Nettekoven et al. 2023). Convergent findings reveal that the regions at and around the Crus I / II lobules respond to higher-order cognitive demands and further that there are multiple, distinct regions embedded within those zones. For example, in a recent comprehensive assessment of functional domains, King et al. (2019) demonstrated robust responses in Crus I / II in an averaged group of participants, sometimes bilateral and sometimes lateralized, across task contrasts that varied demands on language, social cognition, and cognitive control. The responses overlapped but with shifts in the exact spatial positioning and extent between domains supporting both the hypotheses that Crus I / II responds to cognitive demands and also that there may be further specialization. However, it is difficult to appreciate locally specialized regions in group-averaged data given spatial blurring.

Within-individual precision estimates of cerebellar zones associated with cerebral networks provide additional evidence that there may be fine distinctions between closely juxtaposed regions, but also with variability between individuals (Marek et al. 2018; Xue et al. 2021; see also Marek & Greene 2021). In Xue et al. (2021) we reported extensive analyses of two individuals who were each scanned across 31 separate MRI sessions, providing sufficient signal-to-noise ratio (SNR) properties to map local cerebellar regions. We discovered that five separate networks displayed representation in the Crus I / II region extending into the adjacent lobules HVI anteriorly and HVII / HVIII posteriorly. The regions associated with distinct cerebral networks were often interdigitated, with tight juxtapositions that are difficult to resolve in studies that spatially average over participants. Moreover, the multiple regions were linked to cerebral networks that are proposed to participate in distinct domain-flexible and domain-specialized processing functions (Du et al. 2023).

Motivated by the possibility that closely juxtaposed regions of the cerebellum might participate in multiple, distinct cognitive processes, we performed a series of analyses to comprehensively map the human cerebellum in 15 individuals. We then examined the functional response properties within the identified regions using a battery of tasks designed to target multiple facets of higher-order cognition.

## Results

### The Cerebellum Possesses Diverse Higher-Order Cognitive Zones that are Distinct from Sensorimotor Zones

The cerebellum was comprehensively mapped in 15 participants. The full parcellations for two representative participants are displayed in Fig. 1 (P2 and P7; see Supplemental Materials for all other participants). While the exact spatial positions and extent of network representations varied across individuals, the general organization was conserved. A large portion of the anterior lobe of the cerebellum and a smaller subdivision of the posterior lobe were associated with sensorimotor networks (SMOT-A and SMOT-B). Between these somatomotor zones was a large zone populated by regions associated with non-motor networks.

**Figure 1.**
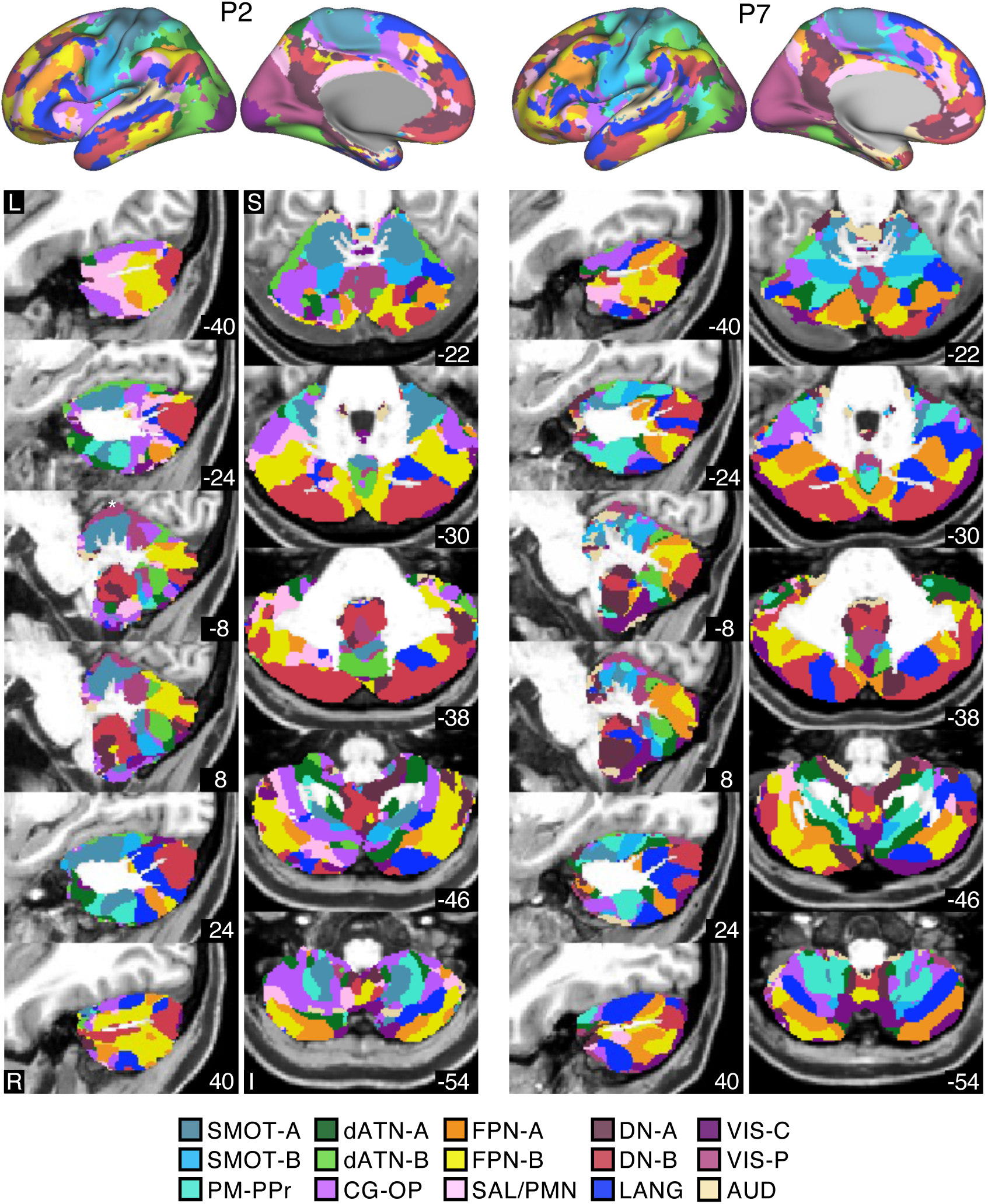
Estimated cerebellar parcellations are displayed for representative participants. (Top) The 15-network Multisession Hierarchical Bayesian Model (MS-HBM) cerebral estimates are displayed for P2 and P7 on the surface. Each network is represented with a different color, labeled at the bottom: Somatomotor-A (SMOT-A), Somatomotor-B (SMOT-B), Premotor-Posterior Parietal Rostral (PM-PPr), Cingulo-Opercular (CG-OP), Salience / Parietal Memory Network (SAL / PMN), Dorsal Attention-A (dATN-A), Dorsal Attention-B (dATN-B), Frontoparietal Network-A (FPN-A)^2^, Frontoparietal Network-B (FPN-B), Default Network-A (DN-A), Default Network-B (DN-B), Language (LANG), Visual-Central (VIS-C), Visual-Peripheral (VIS-P), and Auditory (AUD). **(Bottom)** The cerebellar parcellations associated with the 15 cerebral networks are displayed in sagittal (left) and axial (right) views. No spatial assumptions were made in assigning the cerebellar voxels to cerebral networks. Cerebellar regions linked to the somatomotor networks in the anterior and posterior lobes bound an extensive zone associated with higher-order networks that lies between. The organization is largely symmetrical with at least one exception that the regions linked to the LANG network are more prominent in the right hemisphere. Note the VIS-P ‘strip’ at the superior border (white asterisk) of the cerebellum is likely due to the proximity to adjacent visual cerebral cortex. A permissive cerebellar mask was employed to visualize the full extent of the cerebellar assignments. L, left; R, right; A, anterior; P, posterior; S, superior; and I, inferior. Coordinates in each panel indicate the section level in the MNI152 atlas.

Relevant to the present emphasis on cognitive domains, the apex of the cerebellum near to the horizontal fissure was consistently populated by a megacluster of five higher-order association networks: FPN-A, FPN-B, LANG, DN-B, and DN-A, replicating Xue et al. (2021) and paralleling the motif exhibited in multiple locations across the cerebral cortex (Du et al. 2023) and the striatum (Kosakowski et al. 2023). As a grouping, the megaclusters of five networks accounted for the vast extent of the Crus I and Crus II lobules extending into parts of the adjacent lobule HVI and lobule HVIIb. A cluster of high-order association networks was also observed posteriorly in lobule IX (Fig. 1, axial slice -38). This smaller lobule IX representation was reliably found across participants distant from the larger association megacluster.

To illustrate the consistency of the large Crus I / II megaclusters, cerebellar surface projections are displayed for all 15 participants in Fig. 2. Note that the network estimation procedure did not bias any particular cerebellar location to link to any specific cerebral network, as there were no spatial priors. Yet, in each individual the five networks occupied the same general vicinity of the Crus I / II extended zone. Specifically, the central zone of the megacluster was typically occupied by a large DN-B representation, with a smaller DN-A representation located medially and inferiorly. The representations of DN-A and DN-B were often bounded superiorly and inferiorly by LANG network representations. The LANG network in most, but not all, participants possessed a greater extent in the right cerebellar hemisphere than the left. Some individuals exhibited an especially tight interdigitation of LANG and DN-B1 (Fig. 2; P3, P4, P7, P10). FPN-A and FPN-B bounded the LANG and DN-A / DN-B representations medially and laterally. The organization of the representations of FPN-A and FPN-B was less distinct, with FPN-B displaying a larger representation in most participants. Lateral to the FPN-A and FPN-B network representations, a disjointed set of representations of DN-B (and sometimes DN-A) was observed with a complex and spatially varied pattern of interdigitation across participants.

**Figure 2.**
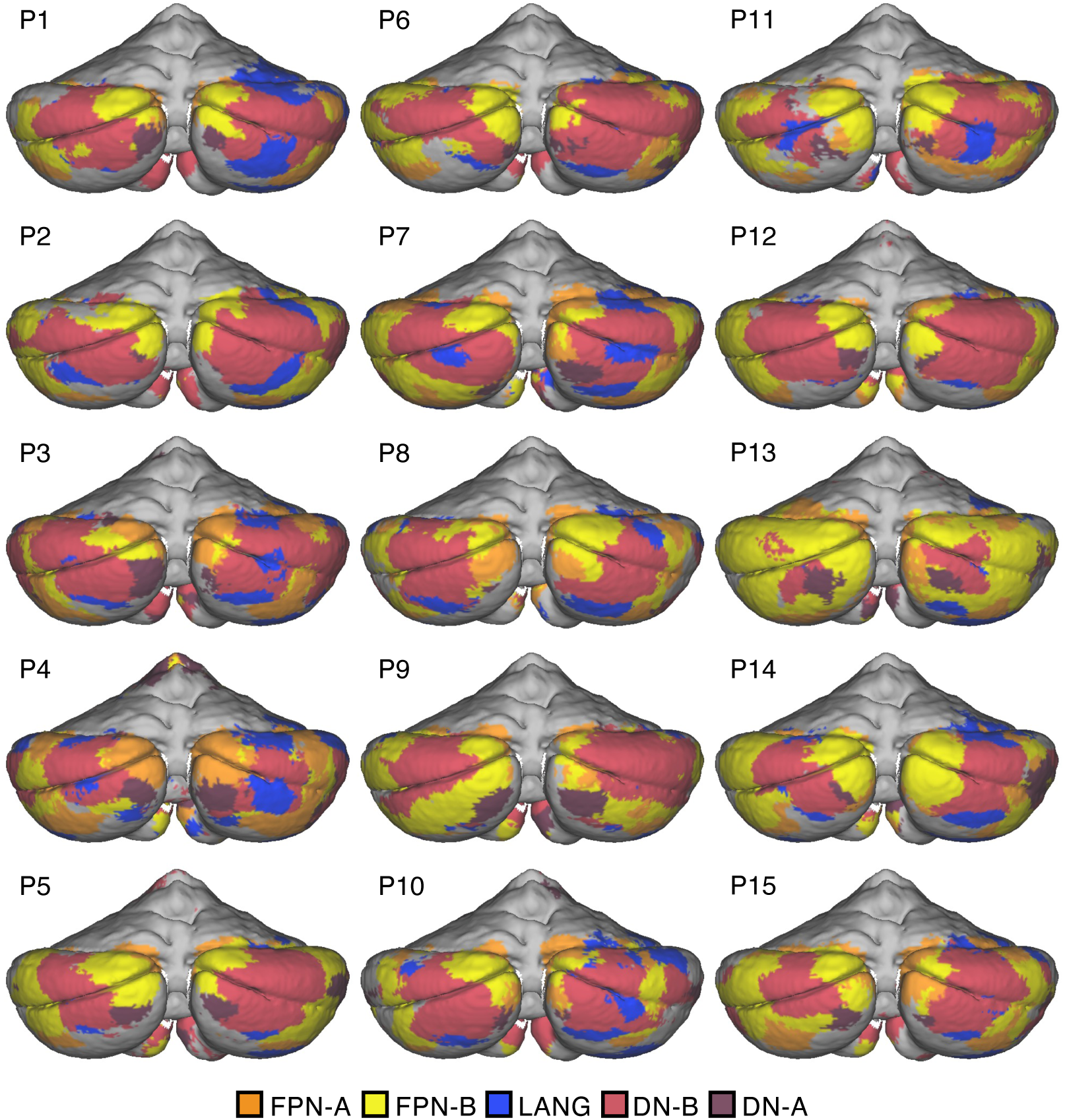
The Crus I / II extended zone is consistently populated by regions linked to five higher-order association networks. Posterior views of the inflated cerebellar surfaces from all 15 participants display regions linked to Frontoparietal Network-A (FPN-A, orange), Frontoparietal Network-A (FPN-B, yellow), Language (LANG, blue), Default Network-B (DN-B, red), and Default Network-A (DN-A, dark red). Across all individuals, the higher-order association regions cover the Crus I / II lobules and extend into the adjacent lobules. Cerebral networks are often linked to multiple cerebellar regions, closely juxtaposed with other network regions creating an interdigitated topography. The exact spatial positioning and extent of network representations varies across individuals, but the general organization is conserved. Notably, the apex of Crus I and Crus II near the horizontal fissure is primarily occupied by regions linked to DN-B, bounded by FPN-A and FPN-B medially and laterally, and by regions linked to the LANG network both anteriorly and posteriorly.

### Closely Juxtaposed Cerebellar Regions are Associated with Distinct Higher-Order Association Networks

Tightly interdigitated, spatially idiosyncratic representations of the distinct association networks were evident in each individual’s cerebellum. This pattern is intriguing but raises the question of whether there is true biological spatial specificity within the cerebellum or, alternatively, whether the tight interdigitation is an artifact of voxels being coupled to multiple networks and the parcellation forcing a winner-takes-all assignment. To address this question, seed-region based functional connectivity was used to explore the specificity of the cerebellar regions without any model assumptions.

For each participant, seed regions were placed within the five different network representations of the Crus I / II extended zone. Figs. 3 and 4 display the cerebral functional connectivity patterns associated with each of these cerebellar seed regions in two representative participants (see Supplemental Materials for all other participants). For each seed region, there was agreement with the individual-specific network boundaries in the cerebral cortex despite the seed-based correlations being free from model constraints, indicating that closely juxtaposed regions of the cerebellum are truly correlated with distinct, spatially specific cerebral networks.

**Figure 3.**
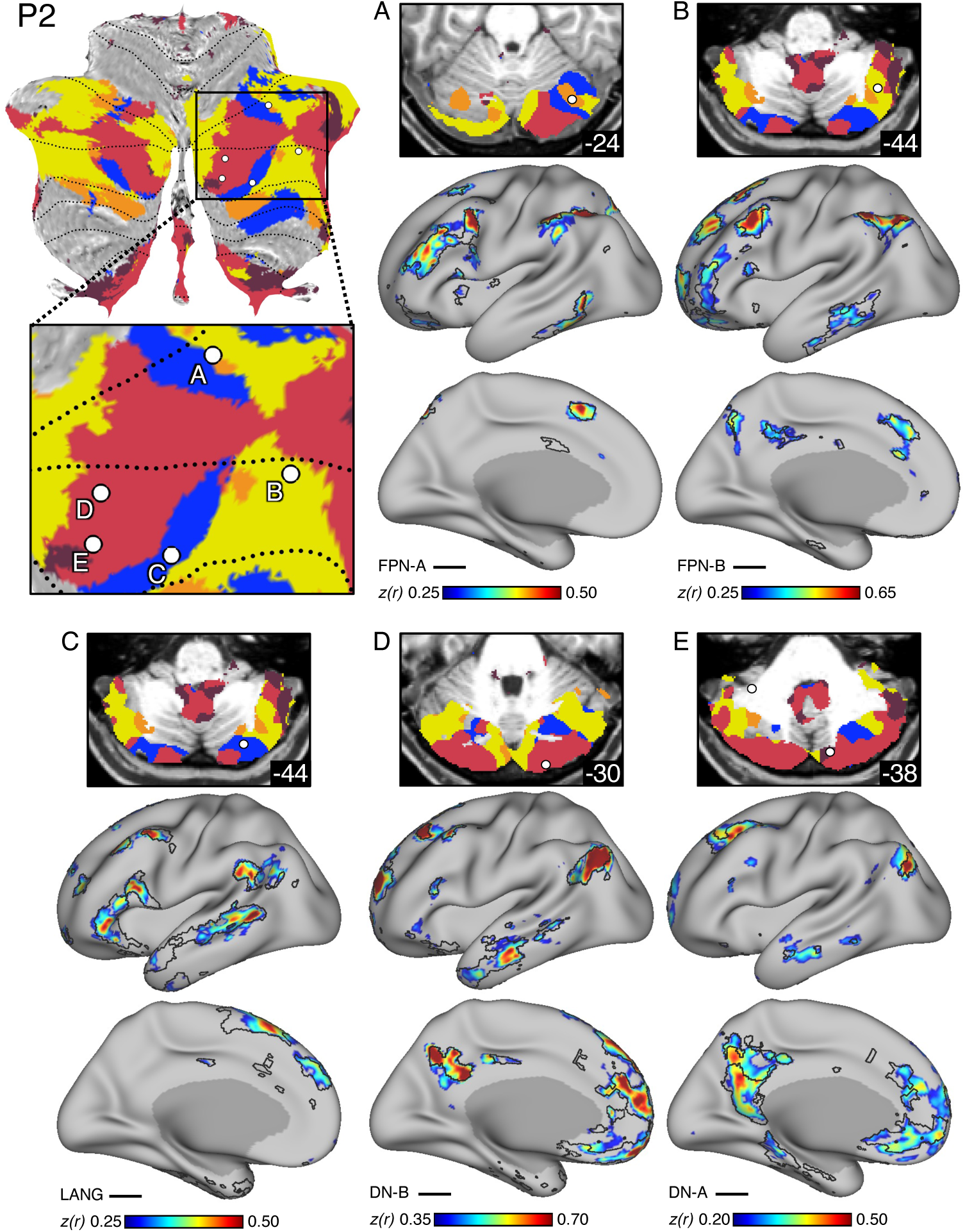
Interdigitated association regions within the Crus I / II extended zone from a representative participant demonstrate anatomical specificity. Correlation patterns in the cerebral cortex are displayed from five distinct cerebellar seed regions of P2 linked to FPN-A **(A)**, FPN-B **(B)**, LANG **(C**), DN-B **(D)**, and DN-A **(E)**. **(Top Left)** To facilitate visualization of the spatial relations among seed regions, the cerebellar parcellation is projected onto a flatmap. Each single-voxel seed region is shown as a white circle on top of the five networks overlayed on the surface. The inset enlarges the relevant portion of the cerebellar flatmap. **(Panels A-E)** The exact location of each seed region is shown on top in the volume. Below is the functional connectivity pattern for that seed region within the left (contralateral) cerebral hemisphere. The color map displays Fisher transformed *r* values. Black lines indicate boundaries of the corresponding within-individual MS-HBM cerebral network estimate. Each seed region correlation pattern recapitulates the full, distributed extent of the relevant network in the cerebral cortex, illustrating specificity of the regions within the Crus I / II extended zone. Note that the DN-A seed region **(E)** projected onto the flatmap gives the illusion of falling within a DN-B zone (most of the depth perpendicular to the surface is assigned to DN-B). As can be visualized in the volume representation, the seed region, which falls deep within the cerebellum, lies within a region assigned to DN-A (for details see the Supplemental Materials). Coordinates in each panel indicate the section level in the MNI152 atlas.

**Figure 4.**
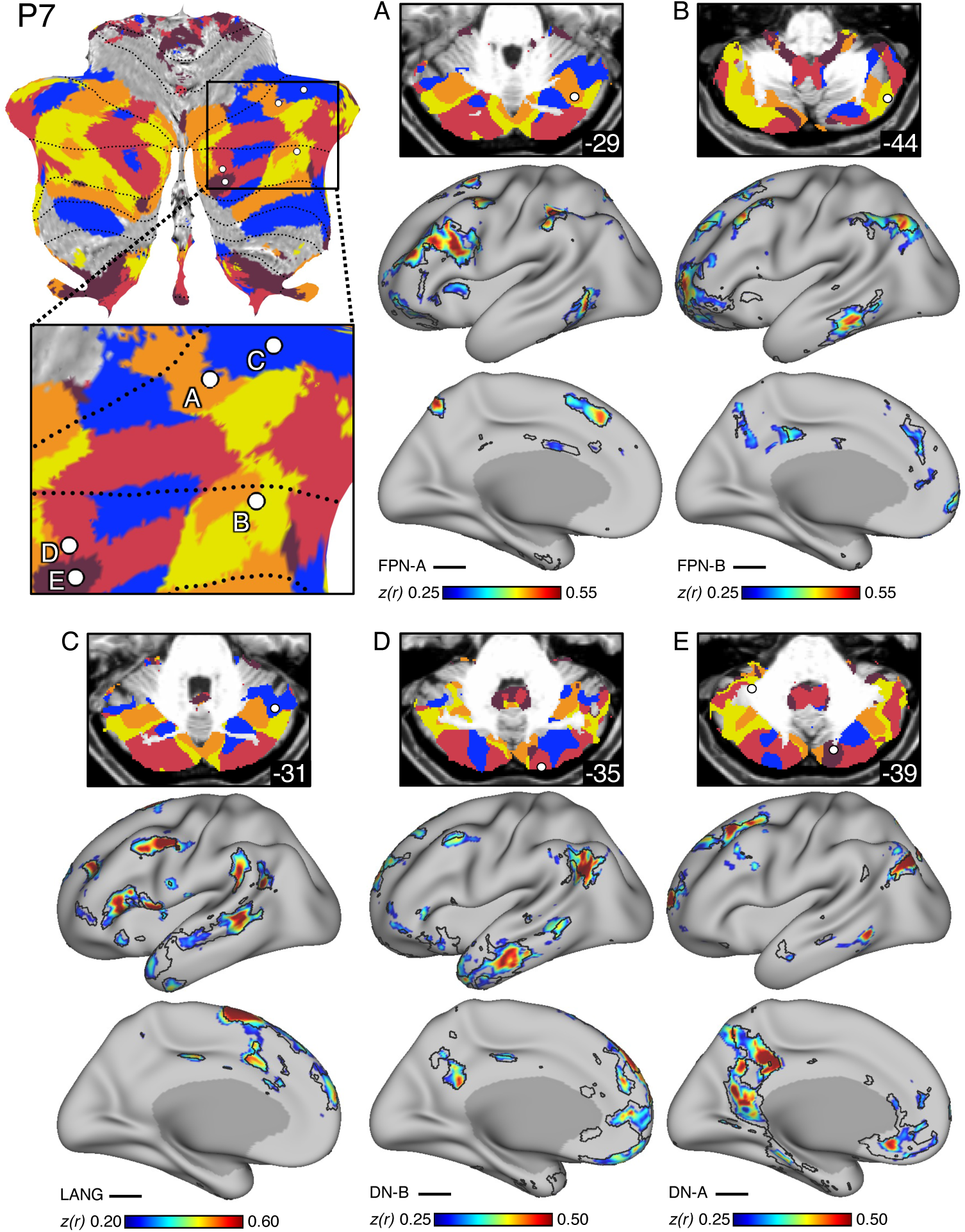
Interdigitated association regions within the Crus I / II extended zone from another representative participant demonstrate anatomical specificity. Paralleling Fig. 3, correlation patterns in the cerebral cortex are displayed from five distinct cerebellar seed regions of P7 linked to FPN-A **(A)**, FPN-B **(B)**, LANG **(C)**, DN-B **(D)**, and DN-A **(E)**. **(Top Left)** The cerebellar parcellation is projected onto a flatmap. Each single-voxel seed region is shown as a white circle on top of the five networks overlayed on the surface. The inset enlarges the relevant portion of the cerebellum. **(Panels A-E)** The exact location of each seed region is shown on top in the volume. Below is the functional connectivity pattern for that seed region within the left (contralateral) cerebral hemisphere. The color map displays Fisher transformed *r* values. Black lines indicate boundaries of the corresponding within-individual MS-HBM cerebral network estimate. The spatial positioning and extent of the networks are different from those of P2, yet each seed region again recapitulates the full, distributed extent of the relevant cerebral network, illustrating specificity of the association networks in the Crus I / II extended zone. Coordinates in each panel indicate the section level in the MNI152 atlas.

For example, the functional connectivity pattern for the seed region placed within the small cerebellar representation of P2’s DN-A was distributed across multiple zones of the cerebral cortex within the boundaries of the DN-A estimated network (Fig. 3E), and less so within the boundaries of the juxtaposed DN-B network (Fig. 3D). Specificity was clear in most participants, following the idiosyncratic anatomical patterns of each individual. To aid visualization of the spatial relations between separate representations in the cerebellum, the five networks and five seed regions were also projected onto the cerebellar flatmap (Figs. 3 and 4; top left).

While our analyses focus on the Crus I / II extended zone, the five higher-order association networks also possessed representations within and near lobule IX (Fig. 1). Lobule IX is located deep within the posterior lobe, beneath the flocculonodular lobe. This representation is spatially distant from the Crus I / II extended zone. DN-A and DN-B showed the most prominent and consistent representation across all 15 individuals. Seed-region based functional connectivity confirmed the specificity of lobule IX representations to the higher-order cerebral association networks in the full set of 15 individuals (see Supplemental Materials). We focus further analyses on the spatial and functional specialization of the Crus I / II extended zone given its larger size and clear pattern in all participants.

### Spatially Disjointed Regions in the Crus I / II Extended Zone Associate with the Same Higher-Order Association Network

Precision estimates within the individual allow for fine details of cerebellar organization to be resolved. We repeatedly observed spatially disjointed regions within each individual’s Crus I / II extended zone that were associated with the same network and separated by regions linked to other association networks. To further explore this observation, we placed seed regions in spatially disjointed representations of the LANG network within the Crus I / II extended zone. We elected to focus on the LANG network because its representation in the cerebellum is particularly disjointed and variable across individuals. Multiple seed regions were placed in distinct LANG representations in the right Crus I / II extended zone of P2 (Fig. 5) and P7 (Fig. 6). Each seed region’s correlation pattern yielded a complete map of the distributed LANG cerebral network despite often being separated in the cerebellum by regions linked to other distinct cerebral association networks.

**Figure 5.**
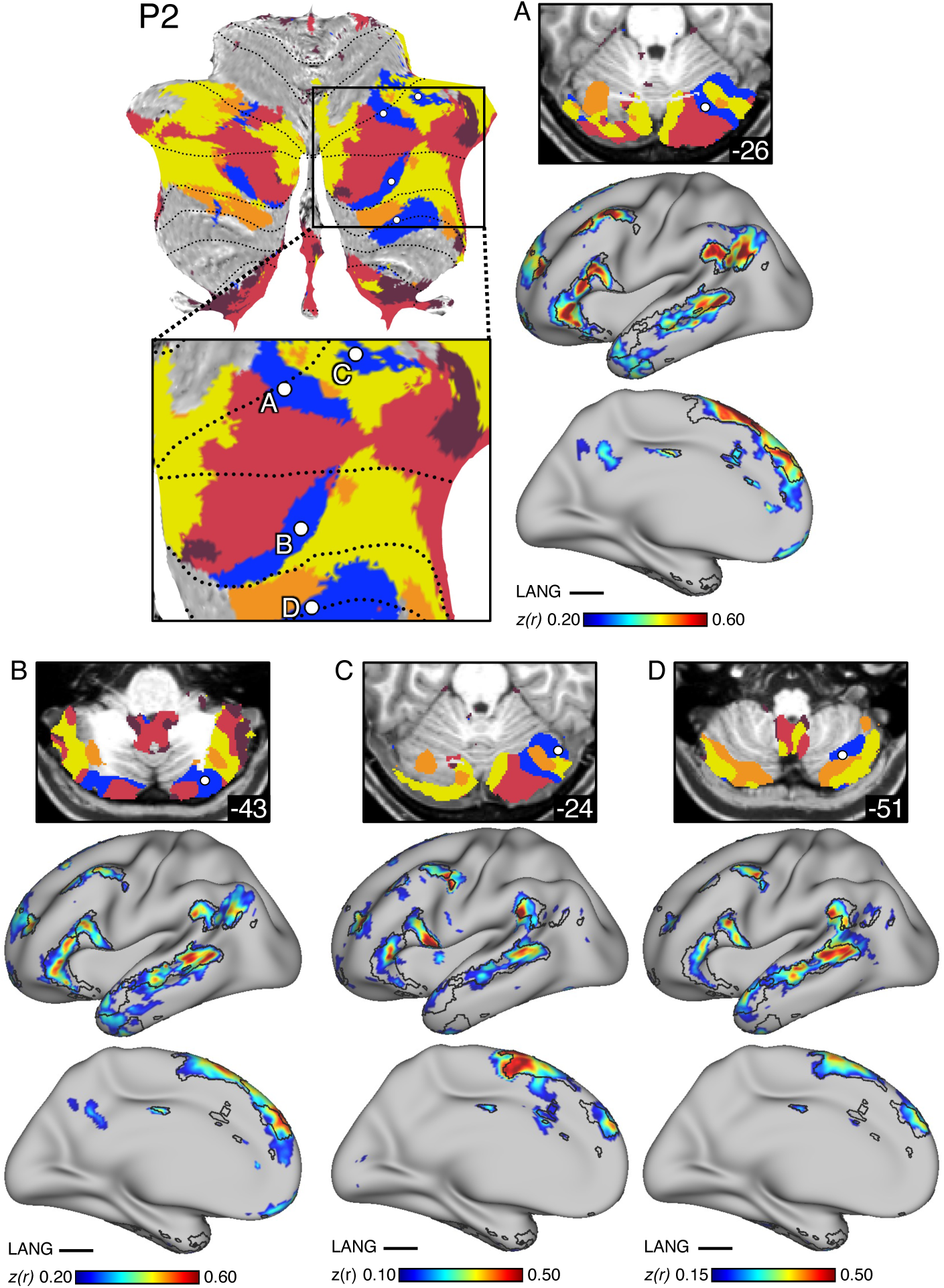
Multiple discontinuous regions in the Crus I / II extended zone are linked to the language network in a representative participant. Correlation patterns in the cerebral cortex are displayed for multiple distinct right cerebellar seed regions of P2 placed within spatially discontinuous representations of the LANG network. **(Top Left)** The cerebellar parcellation is projected onto a flatmap. Each single-voxel seed region is shown as a white circle with the five networks displayed underneath. The inset enlarges the relevant portion of the cerebellum. **(Panels A-D)** The exact location of each seed region is shown on top in the volume. Below is the functional connectivity pattern for that seed region within the left (contralateral) cerebral hemisphere. The color map displays Fisher transformed *r* values. Black lines indicate boundaries of the within-individual MS-HBM cerebral LANG network estimate. Despite being spatially discontinuous, all four seed regions show high specificity (C and D more than A and B) to the cerebral LANG network, across its full distributed extent. Coordinates in each panel indicate the section level in the MNI152 atlas.

**Figure 6.**
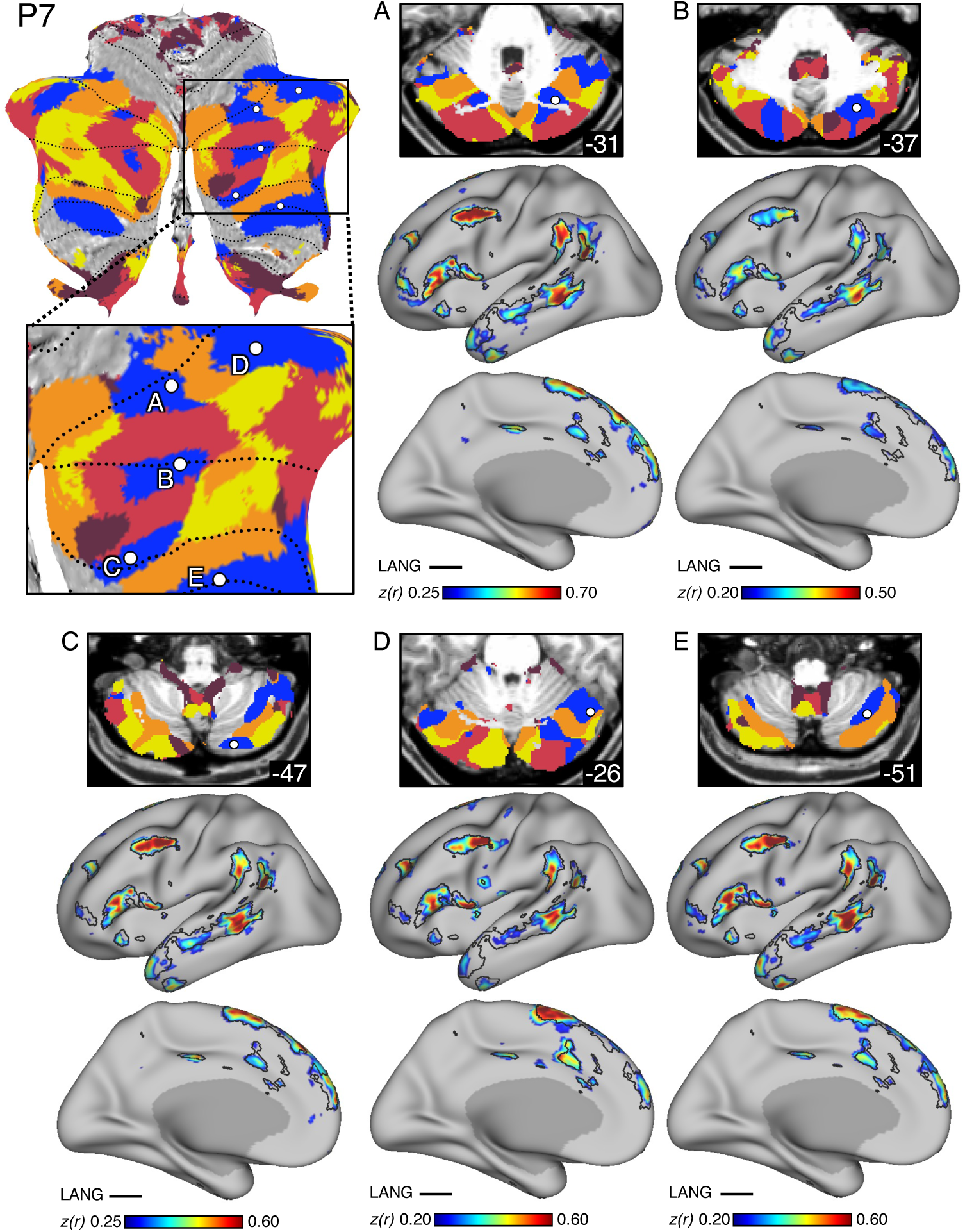
Multiple discontinuous regions in the Crus I / II extended zone are linked to the language network in another representative participant. Paralleling Fig. 5, correlation patterns in the cerebral cortex are displayed for multiple distinct right cerebellar seed regions of P7 placed within spatially discontinuous representations of the LANG network. **(Top Left)** The cerebellar parcellation is projected onto a flatmap using the SUIT toolbox. Each single-voxel seed region is shown as a white circle with the five networks underneath. The inset enlarges the relevant portion of the cerebellum. **(Panels A-E)** The exact location of each seed region is shown on top in the volume. Below is the functional connectivity pattern for that seed region within the left (contralateral) cerebral hemisphere. The color map displays Fisher transformed *r* values. Black lines indicate boundaries of the within-individual cerebral LANG network estimate. Note that seed regions A, C, D, and E are located similarly to P2 regions while the location of seed region B is different. All five spatially separate seed regions again show high specificity to the LANG network, across its full distributed extent. Coordinates in each panel indicate the section level in the MNI152 atlas.

### Regions in the Crus I / II Extended Zone Respond to Domain-Flexible Cognitive Control

The results above suggest that side-by-side regions within the cerebellum are linked to distinct higher-order association networks forming a megacluster of regions in the Crus I / II extended zone. The cerebral counterparts possess robust functional specialization (Du et al. 2023). To test for functional specialization within the cerebellum, we explored task-evoked responses across the multiple regions within the megacluster. Based on prior explorations in the cerebral cortex, we hypothesized that (1) domain-flexible cerebellar regions (associated with networks FPN-A and FPN-B) would functionally dissociate from domain-specialized regions linked to networks LANG, DN-B, and DN-A, and further that (2) the cerebellar regions linked to networks LANG, DN-B, and DN-A would exhibit a triple functional dissociation. Put simply, we hypothesized the cerebellar regions would possess functional response patterns aligned to properties of their partner cerebral networks.

First, we tested working memory task demands. Specifically, we measured the response to high (2-Back) versus low (0-Back) working memory load. We found that the mean response in the merged FPN-A / FPN-B cerebellar region for the N-Back Load Effect task contrast was significantly greater than zero (Fig. 7, left; t(14)=13.06, p<0.001; Cohen’s d=3.37) and was also greater than all other regions in the cerebellar megacluster (paired t-test, all p < 0.001). These results indicate that the region associated with FPN-A / FPN-B responds to working memory load and more so than regions associated with LANG, DN-B and DN-A.

**Figure 7.**
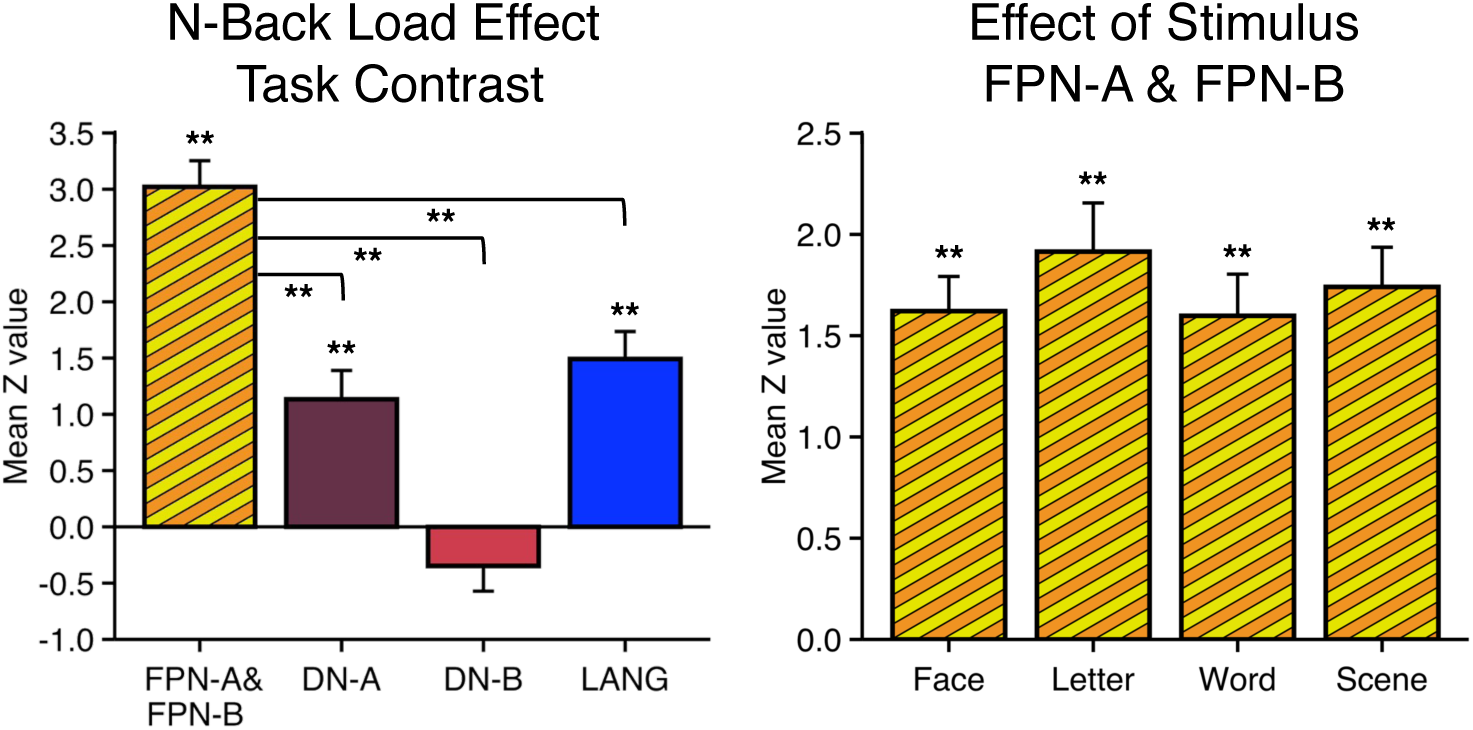
FPN-A and FPN-B regions within the Crus I / II extended zone robustly respond to high working memory load in a domain-flexible manner. Load was manipulated within an N-Back working memory task by contrasting 2-back (high load) versus 0-back (low load) conditions. **(Left)** The bar graph quantifies the mean task contrast *z*-values for the N-Back Load Effect (N = 15) within the *a priori*-defined FPN-A & FPN-B, DN-A, DN-B and LANG regions of the Crus I / II extended zone. The combined cerebellar FPN-A and FPN-B region shows a strong significant positive response that is greater than the responses within the DN-A, DN-B, and LANG regions. **(Right)** The bar graph illustrates the N-Back Load Effect quantified separately for each stimulus domain: face, letter, word and scene. A strong and similar response is observed across all four stimulus domains. Error bars indicate the standard error of the mean. Asterisks indicate significance: ** p <0.001.

Next, we sought to determine whether the response within the FPN-A / FPN-B regions generalized across material domains – a property expected of brain networks that participate in domain-flexible cognitive control (Duncan & Owen 2000; Fedorenko et al. 2013). The N-Back paradigm included four different, verbal and non-verbal, stimulus conditions: Face, Letter, Word and Scene. The mean response for the N-Back Load Effect task contrast within each stimulus type was estimated separately. For the merged FPN-A / FPN-B region, responses to all stimulus types were robust and significantly greater than zero (all p<0.001; Fig. 7, right).

Taken together, these findings suggest that the FPN-A / FPN-B region of the Crus I / II extended zone exhibits a robust domain-flexible response to working memory demands, distinguishing it from the adjacent regions associated with the LANG, DN-B, and DN-A networks.

### Distinct Regions in the Crus I / II Extended Zone Respond to Language, Social, and Spatial / Episodic Task Demands

Cerebellar regions linked to FPN-A / FPN-B responded to working memory demands in a domain-flexible manner. Here, we tested the functional properties of the distinct juxtaposed regions in the Crus I / II extended zone. Specifically, we sought to determine if the cerebellar regions associated with LANG, DN-B and DN-A would functionally dissociate across three contrasts that differentially emphasized language, social, and spatial / episodic task demands (Du et al. 2023).

A repeated measures ANOVA on region-level task response revealed a significant 3 x 3 interaction between the effect of task contrast and region (F(4, 48) = 30.53, p < 0.001). Using paired t-tests, we then tested the response to individual task contrasts, with the hypothesis that each cerebellar region’s within-domain response would be significantly greater than either of the other two regions. In other words, we a priori predicted that (1) the response to the language task contrast would be significantly greater in LANG than in DN-A and DN-B, (2) the response to the social task contrast would be significantly greater in DN-B than LANG or DN-A, and (3) the spatial / episodic task contrast would be greater in DN-A than LANG or DN-B. Five of these six planned comparisons were significant. The Episodic Projection task contrast recruited the DN-A region (Fig. 8, left) significantly more than the DN-B (t(12)=9.16, p<0.001) and LANG (t(12)=10.92, p<0.001) regions. The Theory-of-Mind task contrast recruited the DN-B region (Fig. 8, middle) more the DN-A (t(12)=3.54, p<0.05) and LANG (t(12)=8.45, p<0.001) regions. Finally, the response to the Sentence Processing task contrast in the LANG region (Fig. 8, right) was significantly greater than the response in the DN-A region (t(12)=3.27, p<0.05) and greater but not statistically different from the response in the DN-B region (t(12)=1.16, p=0.13).

**Figure 8.**
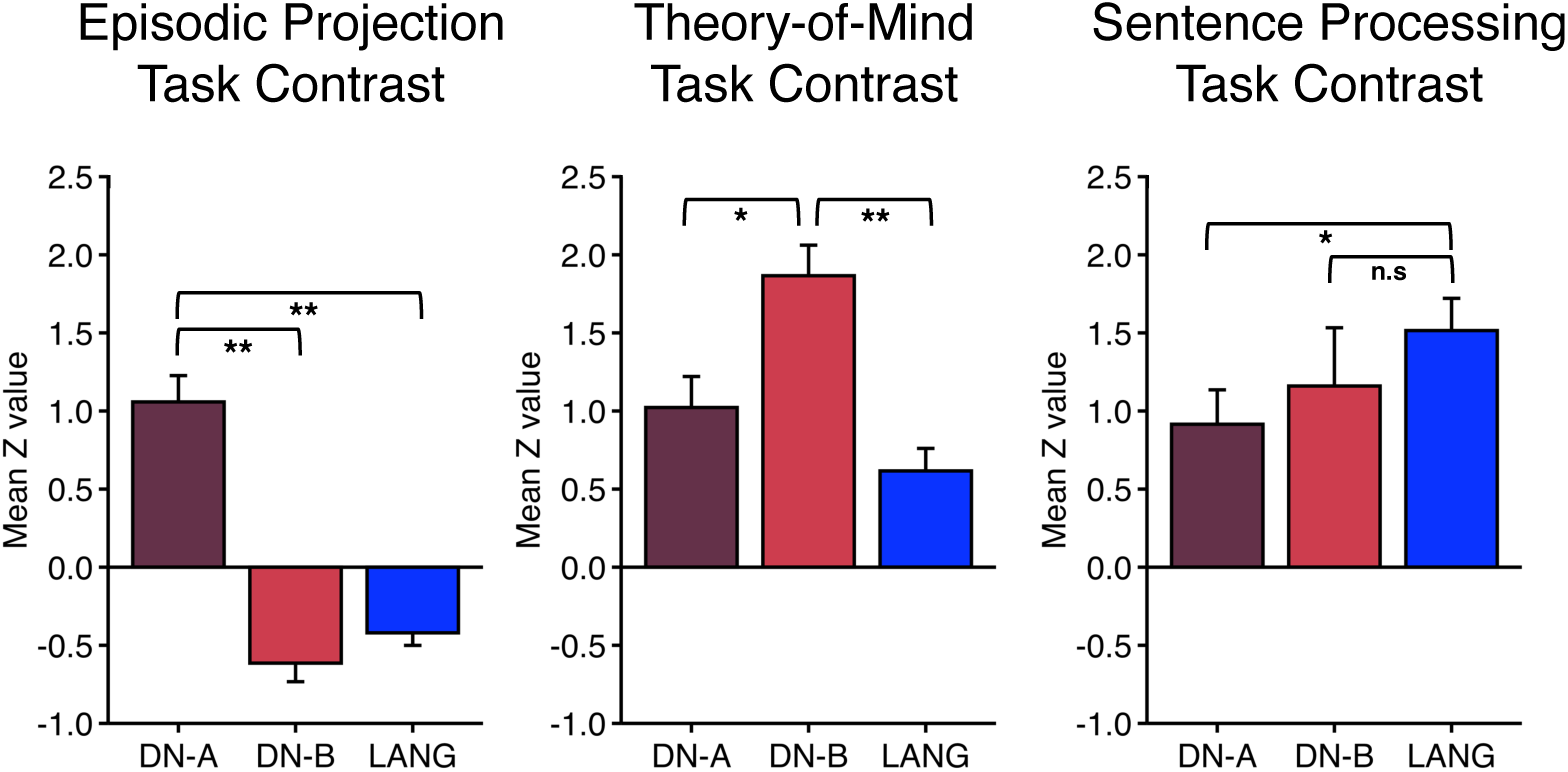
Distinct regions within the Crus I / II extended zone functionally dissociate three higher-order cognitive domains. Distinct domains of higher-order cognitive processing were manipulated across separate task contrasts used previously to explore domain-specificity in the cerebrum. Bar graphs quantify the responses in specific regions of the Crus I / II extended zone to **(Left)** Episodic Projection, **(Middle)** Theory-of-Mind, and **(Right)** Sentence Processing task contrasts. Each panel displays mean *z*-values (N = 13) from the task contrast labelled at top; each bar shows one *a priori*-defined cerebellar region labelled at bottom (DN-A, DN-B, or LANG). The full 3x3 interaction (cerebellar region by task contrast) is significant (*p* < 0.001). Tests of pairwise effects further support the full triple dissociation. The response within the DN-A region is significantly greater than the DN-B and LANG regions in the Episodic Projection task contrast. The response within the DN-B region is significantly greater than the DN-A and LANG regions in the Theory-of-Mind task contrast. The response within the LANG region is significantly greater than the DN-A region and greater (but not significant) than the DN-B region. Error bars indicate the standard error of the mean. Asterisks indicate significance: * p <0.05, ** p <0.001.

These results reveal a triple dissociation among the DN-A, DN-B, and LANG regions within the Crus I/II extended zone, as each region is specialized to support a distinct higher-order cognitive domain.

## Discussion

The present findings suggest that the human cerebellum possesses multiple side-by-side regions specialized for distinct aspects of higher-order cognition. While the spatial positions varied across individuals (Marek et al. 2018; Xue et al. 2021), each individually mapped cerebellum contained a megacluster of regions in the Crus I / II extended zone that was associated with five distinct cerebral association networks (FPN-A, FPN-B, LANG, DN-B and DN-A). The anatomical distinctions among the megacluster regions predicted robust functional dissociations. A trio of domain-specialized regions at the Crus I / II apex of the cerebellum, centered on the horizontal fissure, was surrounded by regions linked to domain-flexible cognitive control, presumably driven by each region’s differential connectivity to its partner cerebral network. Adding a level of complexity to the spatial organization of the cerebellum, the same cerebral network was often associated with multiple discontinuous cerebellar regions interdigitated with regions linked to other networks – a feature that is underappreciated in prior group estimates of functional organization and important to translational goals including neuromodulation.

### Multiple Cerebellar Regions Are Specialized for Distinct Aspects of Cognition

Leiner, Leiner, and Dow (1986; 1989; 1993) introduced the idea that in the large-brained primates, cerebrocerebellar circuits contribute to skilled mental performance in much the same way that distinct circuits contribute to skilled motor performance. They suggested that the cerebellum might best be thought of as a general-purpose computer whose functions “[depend] on the input-output connections that evolved between the cerebellum and other parts of the brain.” (Leiner, Leiner, and Dow 1986). The idea that the cerebellum supports cognitive functions has been amplified by multiple lines of investigation (Schmahmann 2004; Ito 2008; Strick, Dum, & Fiez 2009; Stoodley 2012; Buckner 2013; Sokolov, Miall, & Ivry 2017; Schmahmann et al. 2019; Wagner & Luo 2020). The present results expand our understanding by mapping in detail the complex yet elegant organization of the cognitive domains in the cerebellum.

The Crus I / II extended zone at the apex of the human cerebellum is occupied by multiple regions linked to distinct cerebral association networks. Although there were variations in spatial patterns among individuals, regions associated with at least five networks were consistently identified in each of the 15 individuals examined. This suggests that, despite individual anatomical differences, the existence and spatial organization of the networks exhibit a considerable degree of generality. Prior within-individual explorations of the cerebellum converge (Marek et al. 2018; Xue et al. 2021), and similar megaclusters have been identified in the striatum (Kosakowski et al. 2023). The tight juxtapositions of the multiple regions and their spatial variability between individuals makes it clear why these distinctions were challenging to fully appreciate in past group-based explorations of the cerebellum, including our own (e.g., Buckner et al. 2011). Mapping the regions within each individual’s idiosyncratic anatomy enabled functional distinctions to be probed with a high degree of precision.

Two levels of functional specialization emerged. Suggesting a domain-flexible role in cognitive control, the cerebellar regions linked to the FPN-A and FPN-B networks responded to working memory demands across multiple stimulus domains (Fig. 7). This response profile is characteristic of the multiple-demand network that prominently includes prefrontal regions (Duncan & Owen 2000; Fedorenko, Duncan, & Kanwisher 2013; DiNicola, Sun, & Buckner 2023). By contrast, the cerebellar regions linked to the LANG, DN-B, and DN-A networks displayed differential effects across language, social, and spatial / episodic task domains (Fig. 8). Examining the functional-anatomical details in depth revealed additional insights.

One important anatomical observation is that cerebellar specialization is not described by a linear spatial gradient. The regions linked to FPN-A and FPN-B in the cerebellum tended to surround the domain-specialized regions with multiple, discontinuous regions in each individual. There was not a simple medial to lateral organization, and the spatial variability was such that the approximate anatomical location in one individual linked to domain-flexible cognitive control might be the center of a domain-specialized region in another person. Despite this spatial variability, when the cerebellar regions linked to FPN-A and FPN-B were identified in an individualized manner, they could be demonstrated to robustly modulate to working memory demands to a greater degree than their adjacent domain-preferential regions.

Differential specialization was robust for the domain-preferential cerebellar regions. A particularly informative finding was that the regions associated with DN-B and DN-A revealed a functional double-dissociation between tasks involving making social inferences (Theory-of-Mind) and tasks involving remembering and imagining future scenarios (Episodic Projection). The cerebellar regions specialized for processing in the social domain (Van Overwalle et al. 2020) could be further separated from juxtaposed language regions and from juxtaposed regions supporting cognitive control. The spatial details again varied between individuals as well as between the hemispheres within the same individual. In a revealing observation about the complexity of localization within the cerebellum, the LANG network was associated in some individuals with four or more discontinuous regions with multiple separate adjacencies to the regions linked to the network that responded when making social inferences (Fig. 2).

While the origins of why and how the brain specializes across these domains is an unsolved puzzle, the present evidence provides strong support that the functional specialization of networks prominent in the cerebrum is also present in the cerebellum with complex interdigitated juxtapositions reminiscent of the higher-order association zones of the cerebral cortex.

### Limitations

Employing tasks designed to activate specific cerebral networks proved to be an effective approach to functionally characterize the Crus I / II extended zone. However, it is important to acknowledge a bias inherent in this strategy. Task selection was guided by known functional attributes of the networks in the cerebral cortex. Consequently, this cerebral-centric approach could potentially miss the specific role of the cerebellum in higher-order processing. One clue about divergence might be the equivocal results of the Sentence Processing task contrast (Fig. 8, right). It is not possible from the present data to resolve why that particular dissociation is less robust here in the cerebellum than has been found for the cerebrum (Fedorenko et al. 2012; Braga et al. 2020; Du et al. 2023). Spatial blur and noise associated with technical aspects of imaging the cerebellum could be the origin, but it is also worthwhile to note that the Sentence Processing task involved passive presentation of word strings with an orthogonal participant-directed response at the end of the string to indicate compliance. The response did not specifically refer to the word stimuli and there was no possible right or wrong response. This task detail may be relevant to understand the embedded processing demands that drive the cerebellar response. Future work with a greater range of task manipulations will be required to provide insight.

Another limitation relates to the challenge of visualizing the cerebellar cortex. The complex and detailed spatial arrangements of the cerebellar folia (Sereno et al. 2020) are below the resolution of our experiment, and thus our projections to a model surface are necessarily imperfect approximations of the anatomy (Diedrichsen and Zotow 2015). While the flat and inflated surface representations are valuable tools for orientation, given the spatial limitations of the data we acquired, they likely distort and blur over details. We provide maps in the volume as open resources for the community for further exploration. In the future, explorations of finer spatial details will benefit from higher-field, higher-resolution functional neuroimaging.

### Conclusions

The present study examined the organization and function of the cognitive cerebellum within intensively sampled individual participants. The Crus I / II extended zone is occupied by multiple distinct regions that likely gain their functional properties via their partnered networks in the cerebrum.

## Methods

### Participants

Fifteen adults aged 18–34 yr volunteered for monetary compensation (mean = 22.1 yr, SD = 3.9 yr, 9 women, all right-handed). All were native English speakers with no history of neurological or psychiatric illness, and they came from diverse backgrounds (9 of the 15 individuals self-reported as non-white and / or Hispanic). Informed consent used a protocol approved by the Institutional Review Board of Harvard University. The data were previously reported in a paper focused on the cerebral cortex (Du et al. 2023). Here the data were reanalyzed with a focus on the cerebellum. Specifically, participants were scanned across 8-11 MRI sessions. Three participants also took part in a study of motor movement that included data utilized here (Saadon-Grosman et al. 2022). Each session included multiple resting-state fixation and task runs (Table S1). Task runs relevant to the present analyses targeted four paradigms as well as fixation, described in detail below.

### MRI Data Acquisition

Scanning was carried out at the Harvard Center for Brain Science using a 3T Siemens MAGNETOM Prismafit MRI scanner and the vendor’s 32-channel phased-array head coil (Siemens Healthcare; Erlangen, Germany). Sessions from Saadon-Grosman et al. (2022) used a 64-channel phased-array head-neck coil. fMRI data from the two coils are nearly indistinguishable and have been combined previously (Du et al. 2023), and they are also combined here. Foam and inflatable padding were used to minimize head movement, and participants viewed a rear-projected display positioned to optimize comfortable viewing. Eyes were video recorded using an Eyelink 1000 Core Plus with a Long-Range Mount (SR Research, Ottawa, Ontario, Canada).

A multiband gradient-echo echo-planar pulse sequence sensitive to blood oxygenation level-dependent (BOLD) contrast, provided by the Center for Magnetic Resonance Research (CMRR) at the University of Minnesota, was used for all functional imaging (2.4 mm isotropic voxels, TR = 1,000 ms, TE = 33.0 ms, flip-angle = 64°, matrix 92 × 92 × 65, FOV = 221 × 221 covering the full cerebrum and cerebellum, 5x multislice acceleration). Two dual gradient-echo B0 field maps were also acquired to correct for susceptibility-induced gradient inhomogeneities with slice spatial resolution matched to the BOLD sequence (TE = 4.45 ms, 6.91 ms, TR = 295 ms, flip angle = 55°). The first two sessions of P12 were acquired in a different FOV (211 × 211) that had no observable impact. A high resolution T1w structural scan was acquired using a multi-echo magnetization prepared rapid acquisition gradient echo (ME-MPRAGE; van der Kouwe et al. 2008) sequence (0.8mm isotropic voxels, TR=2500 ms, TE=1.81, 3.6, 5.39, 7.18 ms, TI=1000 ms, 208 slices, flip angle=8°, matrix=320 x 320 x 208, in-plane 2x GRAPPA acceleration). A matched T2w structural scan was also acquired using a sampling perfection with application-optimized contrasts using different flip angle evolution (SPACE) sequence (0.8 mm isotropic voxels, TR=3200 ms, TE=564 ms, 208 slices, matrix=320 x 320 x 208, in-plane 2x GRAPPA acceleration). Rapid T1w ME-MPRAGE structural scans were obtained at least twice in each participant as backup (TR = 2,200 ms, TE = 1.57, 3.39, 5.21, 7.03 ms, TI = 1,100 ms, flip-angle = 7°, voxel size 1.2 mm, matrix 192 x 192 x 176, in-plane 4x GRAPPA acceleration).

### Task Paradigms

fMRI data were acquired during a continuous fixation task to be used for network estimation, and separately during task runs to probe functional specialization. Alertness was monitored during each functional run using eye monitoring. The behavioral tasks are described briefly below and in detail, including exclusions, in Du et al. (2023).

#### Resting-State Fixation

For resting-state fixation runs, participants fixated a centrally presented black crosshair on a light gray background and were instructed to remain still, and awake. Each participant completed 15 to 24 usable runs. Each resting-state fixation run lasted 7 min 2 sec (422 frames with the first 12 frames removed for T1 equilibrium).

#### Working Memory (N-Back) Task

We used an N-Back task (2-back versus 0-back) to target working memory load. Load was manipulated by having 2-back and 0-back conditions in separate blocks. Inspired by Barch et al. (2013), four different stimulus types were included to examine whether the load effect was domain-flexible or domain-specific (Face, Word, Scene, and Letter; see also DiNicola, Sun, & Buckner 2023). Each stimulus type was presented in a separate block. For Face, Scene, and Letter conditions, participants indicated if the current stimulus matched a reference template (0-Back or low load condition) or if it matched the stimulus presented two positions back in the sequence (2-Back or high load condition). For Face, Scene, and Letter conditions, matches were exact repeats of the target. For the Word condition, participants indicated if the current word rhymed with the target word (e.g., “dream” would match “steam”). Each participant completed 8 runs. Each N-Back Task run lasted 4 min 44 sec (284 frames with the first 12 frames removed for T1 equilibrium). The primary comparison of interest was the contrast between 2-back blocks and the 0-back blocks. Blocks with different stimulus types allowed the generality of the effect to be explored.

#### Sentence Processing Task

A sentence processing task adapted from Fedorenko et al. (2010, 2012) targeted processes related to accessing word and phrase-level meaning. Participants passively read real sentences (“IN THE EVENING THE BOY-SCOUTS MADE A FIRE AND BAKED SOME POTATOES”) or pronounceable non-word strings (“ATOMILORM IPTOR CRE FOLE VITE CRE DILES PAME EXFULTER CHILK GOR JESSER”) as a reference control. At the end of each string participants pressed a key to indicate the trial was complete. Word strings and non-word strings were presented in blocks of three strings. Extended fixation blocks were interspersed with task blocks. Six runs were collected for each participant. Each run lasted 5 min and 0 sec (300 frames with the first 12 frames removed for T1 equilibrium). The primary comparison of interest was the contrast between sentence blocks and non-word blocks.

#### Theory-of-Mind Tasks

The Theory-of-Mind (ToM) tasks investigated processes related to understanding other people’s mental states. Adapted from Saxe and colleagues, two paradigms were employed: the False Belief and Pain paradigms (Saxe & Kanwisher 2003; Dodell-Feder et al. 2011; Bruneau et al. 2012; Jacoby et al. 2016). In the False Belief paradigm, participants read brief stories depicting a protagonist with a false belief (False Belief condition) or a physical scene (False Photo condition). In the False Belief condition, participants indicated if the protagonist’s beliefs were true or false; in the False Photo condition, participants indicated whether the story was true or false. The contrast of interest was False Belief compared to False Photo. In the Pain paradigm, stories described emotionally painful situations (Emo Pain condition) and were contrasted with control stories involving physical pain (Phys Pain condition). At the end of each story, participants rated the level of emotional / physical pain experienced by the protagonist during the question period. Eight runs were collected from each participant (four of each paradigm type). Each run lasted 5 min and 18 sec (318 frames with the first 12 frames removed for T1 equilibrium). As both paradigms produce similar task activation patterns, they were combined in the primary contrast of ToM versus control stories (See DiNicola et al. 2020).

#### Episodic Projection Task

The Episodic Projection task targeted processes related to remembering the past and imagining the future (prospection). In the target task conditions, participants read scenarios that oriented them to either a past (Past Self) or a future (Future Self) situation, and then answered questions about the scenarios. The control condition asked participants about a present situation (Present Self). Each run lasted 10 min and 17 sec (617 frames with the first 12 frames removed for T1 equilibrium). Ten runs were collected from each participant. The primary comparison of interest was the contrast between the Past and Future Self conditions versus the Present Self control condition.

### Data Processing and Registration

The data were processed using a custom pipeline to align the data within each individual across multiple runs and sessions, preserving anatomical details and minimizing spatial blur (Braga et al. 2019; Du et al. 2023). The resulting output was registered to both the participant’s T1w image (native space) and to the MNI152 standard space (Evans et al. 2012). Multiple matrices were combined for registration. The first matrix corrected for motion (6 degrees of freedom, DOF; MCFLIRT, FSL; Jenkinson et al. 2012). The second was used to field-map-unwarp the BOLD data (FUGUE, FSL). The third registered the field-map-unwarped data to the mean BOLD template (12 DOF; FLIRT, FSL). The fourth registered the mean BOLD template to the native space T1w image (6 DOF using boundary-based registration; Freesurfer). The fifth and final matrix registered the native space T1w image to the MNI152 atlas using nonlinear registration (FNIRT, FSL). The 0.8 mm isotropic T1w image defined each participant’s native space, along with the matched T2w image when calculating pial and white matter boundary surface estimates (“recon-all”, Freesurfer). For one participant (P12), the registration failed with the 0.8 mm T1w image so we used the backup 1.2 mm image.

The native space data were resampled to the fsaverage6 standard cortical surface (trilinear interpolation; 40,962 vertices per hemisphere; Fischl et al. 1999) and then surface-smoothed using a 2 mm full width at half maximum (FWHM) Gaussian kernel. Data in MNI152 standard space were smoothed with a 4 mm FWHM kernel. Whole brain signal regression was applied to all functional data. For resting-state fixation runs only, ventricular and deep cerebral white matter mean signals, six motion parameters, and their temporal derivatives were also regressed (3dTproject, AFNI; Cox 1996). The residual BOLD data were then band-pass filtered at 0.01–0.1 Hz.

### Individual-Specific Parcellation of the Cerebellum

We previously parcellated the cerebral cortex in all of the present participants using a multisession hierarchical Bayesian model (MS-HBM; Du et al. 2023). We used these cerebral parcellations here to estimate the detailed organization of the cerebellum within each individual. First, we calculated functional connectivity between all cerebral vertices and cerebellar voxels. Then each cerebellar voxel was assigned to its most correlated cerebral network (see Buckner et al. 2011; Xue et al. 2021).

Specifically, to compute functional connectivity, we calculated Pearson’s correlation between the time courses of all cerebral cortical vertices and all cerebellar voxels for each resting-state fixation run of each participant. The resulting connectivity values were Fisher z-transformed and averaged across all runs within each participant. The cerebellum was defined by an oversized (generous) cerebellar mask created by dilating a cerebellar mask generated through Freesurfer’s “recon-all” using a disc with a radius of 6 voxels. To assign network labels to cerebellar voxels, the top 400 cerebral cortical vertices with the strongest correlations to the voxel’s fMRI time course were identified (Xue et al. 2021). The network assigned to the cerebellar voxel was determined by the cerebral cortical network that appeared most frequently among these 400 vertices.

One additional step was conducted to filter assignments of cerebellar voxels to noisy cerebral vertices, particularly to the AUD network, which possesses spurious cerebral assignments in the regions of susceptibility artifact in the ventral temporal lobe and parts of the orbitofrontal cortex (Du et al. 2023). A large, permissive mask was manually defined on the cerebral cortex fsaverge6 surface to include the auditory cortex. Then, all correlations to AUD cerebral vertices outside of this mask were zeroed. As a result, the noisy cerebral vertices were not included but the main, major components of the AUD network were still able to exert their influence on assignments (see Supplemental Materials for example).

### Model-Free Seed-Region Based Functional Connectivity Analysis

The cerebellar parcellation was derived from the individual-specific parcellation of the cerebral cortex, estimated using the MS-HBM. The analyses assumed that there is a one-to-one relation between the networks in the cerebral cortex and regions in the cerebellum. To explore the relation between cerebellar and cerebral cortices without strong assumptions, we undertook a second analysis using a model-free seed-region based approach that was free from prior assumptions (adapted from Du et al. 2023).

For these analyses, single-voxel seed regions were placed in the cerebellum and the unconstrained functional connectivity patterns across the cerebral cortex were derived and visualized. A confirmation of the cerebellar network estimation would be established if a cerebellar seed region within a specific subregion reproduced the corresponding network’s pattern in the cerebral cortex. By contrast, if cerebellar correlation maps from seed regions in the cerebellum were non-specific, spanned multiple networks, or fractionated networks, then the assumptions of the analyses would be undermined. This analysis thus served as a control check.

### Visualization in the Volume and on the Cerebellar Surface

The cerebellum within each individual was visualized within their own anatomy in MNI standard space preserving idiosyncratic details (but within the same space as other participants to allow easy comparison) as well as within the cereballar surface of the MNI template (Diedrichsen & Zotow 2015).

For visualization in the volume, the cerebellum was masked with a dilated volume that was sufficiently generous (and edited as needed) to ensure no cerebellar gray matter voxels were excluded. Specifically, the gray and white matter cerebellar masks generated by each individual’s Freesurfer’s “recon-all” output were transformed to the MNI standard space. The pial mask was dilated twice using a 1-voxel radius disc to expand its boundaries. Subsequently, a mild erosion process was applied to smooth the edges of the mask (performed in Matlab). Next, the white matter mask was subtracted from the modified gray matter mask. Finally, voxels were manually added around any edge as needed to ensure the full cerebellar gray matter was included.

For visualization on the cerebellar surface, we projected the parcellation labels onto flat and inflated representations of the cerebellar surface. These representations were developed by Diedrichsen and Zotow (2015) and integrated into the spatially unbiased infra-tentorial template (SUIT) toolbox (http://www.diedrichsenlab.org/imaging/suit_fMRI.htm). The surface projection method uses an approximate surface of the gray and white matter defined on the MNI standard space template (here specifically, normalized using FSL). The vertices on the two surfaces come in pairs. During the projection process, the algorithm samples the parcellation labels along the line connecting the paired vertices and subsequently assigns the surface vertex with the most frequently occurring label. For this reason, while valuable for comprehensive and simplified visualization, the surface projection can sample across cerebellar voxels with multiple assignments and, in some spatial positions, lose information (see Supplemental Materials). For detailed anatomical explorations, relevant sections of the cerebellar volume representation are shown that preserve exact spatial details along with the comprehensive (but subtly distorted) surface projection.

### Within-individual Task Analysis

The goal of the task analyses was to assess the functional response within network-defined subregions of the cerebellum to explore specialization. Functional task data were analyzed using the general linear model (GLM; FEAT, FSL) as previously described in Du et al. (2023) but here extracting the response from cerebellar regions defined by the cerebellar parcellation. In all instances, the regions were defined within individuals solely based on the functional connectivity network estimates. Then, the functional responses were extracted from the regions prospectively. The focus of the analyses was the megacluster of five networks in Crus I / II and adjacent lobules.

For each individual, regions of interest (ROIs) were defined within the Crus I / II extended region for FPN-A, FPN-B, LANG, DN-B, and DN-A. Due to their similar functional profiles in the cerebral cortex (Du et al. 2023), we made the decision (in advance of examining any cerebellar responses) to combine the FPN-A and FPN-B assignments into a single FPN-A / FPN-B region, resulting in four separate subregions defined within each individual participant’s cerebellum. Specifically, an anatomical mask was created for each individual to include Crus I, Crus II, HVI, and HVIIb lobules based on the anatomical atlas of Diedrichsen et al. (2009). To remove noisy voxels from the regions, all voxels with a temporal SNR (tSNR) below 50 were excluded. We further excluded voxels for which confidence in their network label assignment that was less than 0.4. For each voxel, confidence was calculated as: 1 minus the number of vertices (out of 400) of the second most frequent network divided by the number of vertices of the most frequent network (1-(numVertex1stPlaceNet/numVertex2ndPlaceNet)). Removing ambiguous voxels ensured that, for each individual, subregions that were confidently aligned to distinct networks were targeted. Four distinct regions within the cerebellum were generated for each individual participant (FPN-A / FPN-B, LANG, DN-B, and DN-A).

To quantify the functional response within these a priori-defined cerebellar regions, we calculated the average *z*-value of a specific task contrast for all voxels within each region. We then computed the mean *z*-value across runs to obtain a single response value for each region from each participant. Random effects statistics were then employed to examined response properties with each individual participant contributing a single value representing the mean across their region. Mean *z*-values across participants and standard error of the mean across participants were visualized in the bar graphs.

### Software and Statistical Analysis

Cerebellar parcellations were derived from the cerebral cortex parcellation in Du et al. 2023 and were created using code from Xue et al. (2021), available on: https://github.com/ThomasYeoLab/CBIG/tree/master/stable_projects/brain_parcellation/Xue2021_IndCereb <u>ellum</u>. Seed-region based analyses were conducted using the cerebro-cerebellar correlation matrix. The connectivity patterns were visualized interactively, and seed regions were selected using Connectome Workbench v1.3.2. All statistical analyses were performed using R v3.6.2.

## Acknowledgments

S. Kaiser and J. Ladopoulou assisted with data preparation and upload for open release. We thank the Harvard Center for Brain Science neuroimaging core and FAS Division of Research Computing. We thank T. O’Keefe for assisting in optimization of data processing and R. Mair for MRI physics support, and J. Diedrichsen for providing the SUIT toolbox. The multi-band EPI sequence was generously provided by the Center for Magnetic Resonance Research (CMRR) at the University of Minnesota.

## Footnotes

1 Surface approximations of the highly foliated cerebellar cortex are useful to orient and visualize the relations among cerebellar regions (Diedrichsen & Zotow 2015). However, within our data and methods, the surface representation is a simplified view that does not account for the tight folia. Moreover, a winner-takes-all approach assigned a single value to a surface vertex derived from multiple voxels that spanned the depth of the cerebellar cortex. In instances where there were multiple, distinct voxel assignments, information was lost. For this reason, all analyses were conducted in the volume. Projections to the surface were used to appreciate spatial relations but not relied on for quantification.

2 The Frontoparietal Network (FPN) has been fractionated into distinct, parallel networks in multiple prior studies, but has not been consistently named. Here and in the earlier Du et al. (2023) paper on the cerebral cortex we label FPN-A and FPN-B with the order (A/B naming convention) of Kong et al. (2019) and Xue et al. (2021). We caution the reader that other studies have used the reverse convention, which could lead to confusion (e.g., Braga et al. 2020; DiNicola, Ariyo, & Buckner 2023).

## References

1. Allen G, McColl R, Barnard H, Ringe WK, Fleckenstein J, Cullum CM. Magnetic resonance imaging of cerebellar–prefrontal and cerebellar–parietal functional connectivity. NeuroImage 28: 39–48, 2005.

2. Barch DM, Burgess GC, Harms MP, Petersen SE, Schlaggar BL, Corbetta M, Glasser MF, Curtiss S, Dixit S, Feldt C, Nolan D, Bryant E, Hartley T, Footer O, Bjork JM, Poldrack R, Smith S, Johansen-Berg H, Snyder AZ, Van Essen DC. Function in the human connectome: Task-fMRI and individual differences in behavior. NeuroImage 80: 169–189, 2013.

3. Braga RM, Buckner RL. Parallel interdigitated distributed networks within the individual estimated by intrinsic functional connectivity. Neuron 95: 457–471.e5, 2017.

4. Braga RM, DiNicola LM, Becker HC, Buckner RL. Situating the left-lateralized language network in the broader organization of multiple specialized large-scale distributed networks. J Neurophysiol 124: 1415–1448, 2020.

5. Braga RM, Van Dijk KRA, Polimeni JR, Eldaief MC, Buckner RL. Parallel distributed networks resolved at high resolution reveal close juxtaposition of distinct regions. J Neurophysiol 121: 1513–1534, 2019.

6. Bruneau EG, Pluta A, Saxe R. Distinct roles of the ‘Shared Pain’ and ‘Theory of Mind’ networks in processing others’ emotional suffering. Neuropsychologia 50: 219–231, 2012.

7. Buckner RL. The cerebellum and cognitive function: 25 years of insight from anatomy and neuroimaging. Neuron 80: 807–815, 2013.

8. Buckner RL, Krienen FM, Castellanos A, Diaz JC, Yeo BTT. The organization of the human cerebellum estimated by intrinsic functional connectivity. J Neurophysiol 106: 2322–2345, 2011.

9. Cox RW. AFNI: Software for analysis and visualization of functional magnetic resonance neuroimages. Comput Biomed Res 29: 162–173, 1996.

10. Desmond JE, Fiez JA. Neuroimaging studies of the cerebellum: Language, learning and memory. Trends Cogn Sci 2: 355–362, 1998.

11. Diedrichsen J, Balsters JH, Flavell J, Cussans E, Ramnani N. A probabilistic MR atlas of the human cerebellum. NeuroImage 46: 39–46, 2009.

12. Diedrichsen J, Zotow E. Surface-based display of volume-averaged cerebellar imaging data. PLoS One 10: e0133402, 2015.

13. DiNicola LM, Braga RM, Buckner RL. Parallel distributed networks dissociate episodic and social functions within the individual. J Neurophysiol 123: 1144–1179, 2020.

14. DiNicola LM, Ariyo OI, Buckner RL. Functional specialization of parallel distributed networks revealed by analysis of trial-to-trial variation in processing demands. J Neurophysiol 129: 17–40, 2023.

15. DiNicola LM, Sun W, Buckner RL. Side-by-side regions in dorsolateral prefrontal cortex estimated within the individual respond differentially to domain-specific and domain-flexible processes. J Neurophysiol 130: 1602–1615, 2023.

16. Dodell-Feder D, Koster-Hale J, Bedny M, Saxe R. fMRI item analysis in a theory of mind task. NeuroImage 55: 705–712, 2011.

17. Du J, DiNicola LM, Angeli PA, Saadon-Grosman N, Sun W, Kaiser S, Ladopoulou J, Xue A, Yeo BTT, Eldaief MC, Buckner RL. Within-individual organization of the human cerebral cortex: Networks, global topography, and function. bioRxiv 2023.08.08.552437, 2023.

18. Duncan J, Owen AM. Common regions of the human frontal lobe recruited by diverse cognitive demands. Trends Cogn Sci 23: 475–483, 2000.

19. Evans AC, Janke AL, Collins DL, Baillet S. Brain templates and atlases. NeuroImage 62: 911–922, 2012.

20. Fedorenko E, Hsieh P-J, Nieto-Castañón A, Whitfield-Gabrieli S, Kanwisher N. New method for fMRI investigations of language: Defining ROIs functionally in individual subjects. J Neurophysiol 104: 1177– 1194, 2010.

21. Fedorenko E, Duncan J, Kanwisher N. Language-selective and domain-general regions lie side by side within Broca’s area. Curr Biol 22: 2059–2062, 2012.

22. Fedorenko E, Duncan J, Kanwisher N. Broad domain generality in focal regions of frontal and parietal cortex. Proc Natl Acad Sci USA 110: 16616–16621, 2013.

23. Fischl B, Sereno MI, Dale AM. Cortical surface-based analysis: II: Inflation, flattening, and a surface-based coordinate system. NeuroImage 9: 195–207, 1999.

24. Guell X, Gabrieli JDE, Schmahmann JD. Triple representation of language, working memory, social and emotion processing in the cerebellum: Convergent evidence from task and seed-based resting-state fMRI analyses in a single large cohort. NeuroImage 172: 437–449, 2018.

25. Guell X, Schmahmann J. Cerebellar functional anatomy: A didactic summary based on human fMRI evidence. Cerebellum 19: 1–5, 2020.

26. Habas C. Functional connectivity of the cognitive cerebellum. Front Syst Neurosci 15: 27, 2021.

27. Habas C, Kamdar N, Nguyen D, Prater K, Beckmann CF, Menon V, Greicius MD. Distinct cerebellar contributions to intrinsic connectivity networks. J Neurosci 29: 8586–8594, 2009.

28. Ito M. Control of mental activities by internal models in the cerebellum. Nat Rev Neurosci 9: 304–313, 2008.

29. Jacoby N, Bruneau E, Koster-Hale J, Saxe R. Localizing pain matrix and theory of mind networks with both verbal and non-verbal stimuli. NeuroImage 126: 39–48, 2016.

30. Jenkinson M, Beckmann CF, Behrens TEJ, Woolrich MW, Smith SM. FSL. NeuroImage 62: 782–790, 2012.

31. Kelly RM, Strick PL. Cerebellar loops with motor cortex and prefrontal cortex of a nonhuman primate. J Neurosci 23: 8432–8444, 2003.

32. Keren-Happuch E, Chen SA, Ho MR, Desmond JE. A meta-analysis of cerebellar contributions to higher cognition from PET and fMRI studies. Hum Brain Mapp 35: 593–615, 2012.

33. King M, Hernandez-Castillo CR, Poldrack RA, Ivry RB, Diedrichsen J. Functional boundaries in the human cerebellum revealed by a multi-domain task battery. Nat Neurosci 22: 1371–1378, 2019.

34. Kosakowski HL, Saadon-Grosman N, Du J, Eldaief ME, Buckner RL. Human striatal association megaclusters. bioRxiv 2023.10.03.560666, 2023.

35. van der Kouwe AJW, Benner T, Salat DH, Fischl B. Brain morphometry with multiecho MPRAGE. NeuroImage 40: 559–569, 2008.

36. Koziol LF, Budding D, Andreasen N, D’Arrigo S, Bulgheroni S, Imamizu H, Ito M, Manto M, Marvel C, Parker K, Pezzulo G, Ramnani N, Riva D, Schmahmann J, Vandervert L, Yamazaki T. Consensus paper: The cerebellum’s role in movement and cognition. Cerebellum 13: 151–177, 2014.

37. Krienen FM, Buckner RL. Segregated fronto-cerebellar circuits revealed by intrinsic functional connectivity. Cereb Cortex 19: 2485–2497, 2009.

38. Laumann TO, Gordon EM, Adeyemo B, Snyder AZ, Joo SJ, Chen M-Y, Gilmore AW, McDermott KB, Nelson SM, Dosenbach NUF, Schlaggar BL, Mumford JA, Poldrack RA, Petersen SE. Functional system and areal organization of a highly sampled individual human brain. Neuron 87: 657–670, 2015.

39. Leiner HC, Leiner AL, Dow RS. Does the cerebellum contribute to mental skills? Behav Neurosci 100: 443– 454, 1986.

40. Leiner HC, Leiner AL, Dow RS. Reappraising the cerebellum: What does the hindbrain contribute to the forebrain? Behav Neurosci 103: 998–1008, 1989.

41. Leiner HC, Leiner AL, Dow RS. Cognitive and language functions of the human cerebellum. Trends Cogn Sci 16: 444–447, 1993.

42. Lu J, Liu H, Zhang M, Wang D, Cao Y, Ma Q, Rong D, Wang X, Buckner RL, Li K. Focal pontine lesions provide evidence that intrinsic functional connectivity reflects polysynaptic anatomical pathways. J Neurosci 31: 15065–15071, 2011.

43. Marek S, Siegel JS, Gordon EM, Raut RV, Gratton C, Newbold DJ, Ortega M, Laumann TO, Adeyemo B, Miller DB, Zheng A, Lopez KC, Berg JJ, Coalson RS, Nguyen AL, Dierker D, Van AN, Hoyt CR, McDermott KB, Norris SA, Shimony JS, Snyder AZ, Nelson SM, Barch DM, Schlaggar BL, Raichle ME, Petersen SE, Greene DJ, Dosenbach NUF. Spatial and temporal organization of the individual human cerebellum. Neuron 100: 977–993.e7, 2018.

44. Marek S, Greene DJ. Precision functional mapping of the subcortex and cerebellum. Curr Opin Behav Sci 40: 12–18, 2021.

45. Middleton FA, Strick PL. Anatomical evidence for cerebellar and basal ganglia involvement in higher cognitive function. Science 266: 458–461, 1994.

46. Middleton FA, Strick PL. Cerebellar projections to the prefrontal cortex of the primate. J Neurosci 21: 700– 712, 2001.

47. Nettekoven C, Zhi D, Shahshahani L, Pinho AL, Saadon-Grosman N, Diedrichsen J. A hierarchical atlas of the human cerebellum for functional precision mapping. bioRxiv 2023.09.14.557689, 2023.

48. O’Reilly JX, Beckmann CF, Tomassini V, Ramnani N, Johansen-Berg H. Distinct and overlapping functional zones in the cerebellum defined by resting state functional connectivity. Cereb Cortex 20: 953–965, 2010.

49. Petersen SE, Fox PT, Posner MI, Mintun M, Raichle ME. Positron emission tomographic studies of the processing of singe words. J Cogn Neurosci 1: 153–170, 1989.

50. Saadon-Grosman N, Angeli PA, DiNicola LM, Buckner RL. A third somatomotor representation in the human cerebellum. J Neurophysiol 128: 1051–1073, 2022.

51. Saxe R, Kanwisher N. People thinking about thinking people: The role of the temporo-parietal junction in “theory of mind.” NeuroImage 19: 1835–1842, 2003.

52. Schmahmann JD. An emerging concept: The cerebellar contribution to higher function. Arch. Neurol 48: 1178–1187, 1991.

53. Schmahmann JD. Disorders of the cerebellum: Ataxia, dysmetria of thought, and the cerebellar cognitive affective syndrome. J Neuropsychiatry Clin Neurosci 16: 367–378, 2004.

54. Schmahmann JD. The cerebellum and cognition. Neurosci Lett 688: 62–75, 2019.

55. Schmahmann JD, Guell X, Stoodley CJ, Halko MA. The theory and neuroscience of cerebellar cognition. Annu Rev Neurosci 42: 337–364, 2019.

57. Sereno MI, Diedrichsen J, Tachrount M, Testa-Silva G, d’Arceuil H, De Zeeuw C. The human cerebellum has almost 80% of the surface area of the neocortex. Proc Natl Acad Sci U S A 117: 19538–19543, 2020.

58. Sokolov AA, Miall RC, Ivry RB. The cerebellum: Adaptive prediction for movement and cognition. Trends Cogn. Sci 21: 313–332, 2017.

59. Stoodley CJ. The cerebellum and cognition: evidence from functional imaging studies. Cerebellum 11: 352– 365, 2012.

60. Stoodley CJ, Schmahmann JD. Functional topography in the human cerebellum: A meta-analysis of neuroimaging studies. NeuroImage 44: 489–501, 2009.

61. Strick PL, Dum RP, Fiez JA. Cerebellum and nonmotor function. Annu Rev Neurosci 32: 413–434, 2009.

62. Van Overwalle F, Manto M, Cattaneo Z, Clausi S, Ferrari C, Gabrieli JDE, Guell X, Heleven E, Lupo M, Ma Q, Michelutti M, Olivito G, Pu M, Rice LC, Schmahmann JD, Siciliano L, Sokolov AA, Stoodley CJ, van Dun K, Vandervert L, Leggio M. Consensus paper: cerebellum and social cognition. Cerebellum 19: 833–868, 2020.

63. Wagner MJ, Luo L. Neocortex-cerebellum circuits for cognitive processing. Trends Neurosci 43: 42–54, 2020.

64. Xue A, Kong R, Yang Q, Eldaief MC, Angeli PA, DiNicola LM, Braga RM, Buckner RL, Yeo BTT. The detailed organization of the human cerebellum estimated by intrinsic functional connectivity within the individual. J Neurophysiol 125: 358–384, 2021.

